# Plasmids Across Datasets: Resistance, Virulence, Mobility, and Host Taxonomy

**DOI:** 10.1101/2025.01.30.635773

**Authors:** Célia P.F. Domingues, João S. Rebelo, Francisco Dionisio, Teresa Nogueira

**Affiliations:** INIAV—Instituto Nacional de Investigação Agrária e Veterinária, 4485-655 Vairão, Portugal; cE3c—Centre for Ecology, Evolution and Environmental Changes & CHANGE, Global Change and Sustainability Institute, Faculdade de Ciências, Universidade de Lisboa, 1749-016 Lisboa, Portugal

## Abstract

Plasmids are autonomous DNA molecules that can replicate independently and transfer horizontally between bacterial cells. They play a key role in disseminating adaptive traits, such as antimicrobial resistance and virulence. Understanding plasmid mobility and its association with these traits is crucial to microbial ecology, public health and genomic surveillance. Several databases have been developed to catalogue plasmids assembled from bacterial isolates and metagenomic samples. However, differences in database construction and curation can introduce biases that affect subsequent analyses. In this study, we compare three distinct plasmid genome datasets — the NCBI Reference Sequence Database (RefSeq), the Integrated Microbial Genomes & Microbiomes system (IMG/PR) from bacterial isolates (I) and microbiomes (M) — to assess the influence of data origin on inferences about plasmid mobility types, antimicrobial resistance genes (ARGs), virulence genes (VGs) and host taxonomy. Our analysis reveals that plasmids assembled from metagenomes tend to be smaller than those assembled from isolates. RefSeq plasmids are enriched in conjugative plasmids (pCONJ) and display a higher frequency of ARGs and VGs. In contrast, regardless of whether they originate from isolates or metagenomes, IMG/PR plasmids are enriched in mobilizable plasmids (pMOB). Furthermore, ARGs are more frequently associated with highly mobile plasmids, particularly pCONJ. These findings highlight the importance of database selection in studies of plasmid epidemiology, functional potential and mobility. Standardised curation practices and cross-database comparisons are essential to ensure robust and reproducible insights into plasmid-mediated gene flow.

**Importance:** Plasmids are DNA molecules that can replicate and transfer between bacteria, thereby helping to spread genes that enable bacteria to survive and adapt in different environments. This gene exchange plays a significant part in bacterial evolution. Researchers study these processes using plasmid databases, but the way these databases are constructed can influence the conclusions that are drawn. In this study, we found that key traits, such as those involved in antibiotic resistance and the ability to cause disease, are more often linked to plasmids with greater mobility, particularly in databases containing more clinical samples. However, our results demonstrate that the choice of dataset can significantly impact our understanding of the dissemination of critical genes among bacteria. These findings are valuable for tracking and controlling antibiotic resistance and disease, and highlight the need for carefully constructed, representative databases to support accurate research into how bacteria share genes.

## Introduction

Plasmids, extrachromosomal self-replicating genetic elements, are ubiquitous in bacteria. For example, a recent study identified a specific plasmid, pBI143, as the most prevalent and abundant genetic element in human gut microbiomes of industrialized populations worldwide (1). Moreover, studies involving a few thousand strains have shown that there are, on average, two to three different plasmids per strain of *Escherichia coli, Klebsiella* and *Salmonella*, and almost one plasmid per strain of *Staphylococcus aureus* (2, 3). Plasmids have been extensively studied because they are major drivers of horizontal gene transfer with evolutionary and ecological impact (4), very often may harbour genes that facilitate the degradation of environmental pollutants (5), biofilm formation, and confer bacterial virulence, drug and heavy metal resistance (6–8).

Understanding how plasmids spread is crucial for assessing their role in the dissemination of traits such as antibiotic resistance or virulence. To support such studies, researchers rely on plasmid databases, each developed with specific objectives and containing different information. Plasmids can be isolated from known organisms or assembled from complex microbial communities (metagenomically assembled plasmids). Since they originate from diverse sources and hosts, databases vary significantly in the traits they capture. Among these, one particularly important characteristic that directly influences the epidemic potential of plasmids is their ability to transfer between bacterial cells. Conjugative plasmids (pCONJ) can transfer themselves by replication between bacterial cells, often between different strains or species. Therefore, several works have emphasized their pivotal role in bacterial evolution by acquiring clinically significant traits (9–13). pCONJ are self-transmissible because they encode a complete gene set necessary for conjugation, namely a relaxase (MOB), an endonuclease, that cuts one of the DNA strands at the nic site of the respective *oriT*, and the complete machinery for mating pair formation (MPF), which connects donor and recipient cells while serving as a channel for the plasmid transfer (14, 15). Mobilizable plasmids (pMOB) lack part or all MPF genes, so, for transfer, they must hijack these genes’ products from other coresident conjugative plasmids. Plasmids lacking relaxases but containing the *oriT* are also mobilizable and belong to the pMOB group (3). Finally, non-transmissible plasmids (pNT) have no identifiable *oriT* locus or any of the genes mentioned above, suggesting they cannot transfer horizontally. However, these plasmids may yet contain any other unidentified *oriT* (3, 16).

Previous studies estimated that about one-quarter of the plasmids are pCONJ: 23% in a database comprising 11,386 plasmids from *Escherichia coli* and *Staphylococcus aureus*; and 25.2% in a database comprising 14,029 plasmids (10, 17). However, most likely, previously sequenced plasmids are of clinical interest because they were isolated from human pathogens or because they confer drug resistance. In recent years, new and cheaper sequencing techniques have allowed researchers to sequence and study plasmids from different contexts.

This study aims to understand whether different plasmid sources of different databases are differentially enriched regarding resistance genes, virulence genes, plasmid mobility types, and host taxonomy. NCBI Reference Sequence Database (18) and IMG/PR (19) are two of the largest public repositories of plasmid sequences. While RefSeq contains only plasmids from isolate genomes, IMG/PR includes isolate-derived and metagenome-assembled plasmids. As these databases are used to draw meaningful conclusions about plasmid biology, a key question emerges: to what extent might the choice of database lead to different interpretations or findings? This issue is particularly relevant given that these datasets originate from distinct sampling sources. This analysis aims to evaluate whether the plasmid traits differ significantly between datasets and to identify potential biases associated with the source and context of sampling.

## Results

This study aims to elucidate the differences among three plasmid datasets in terms of their mobility, the presence of antimicrobial resistance genes (ARGs) and virulence genes (VGs), as well as host taxonomy. We analyzed three distinct datasets: plasmids identified from RefSeq isolates, plasmids identified from IMG/PR isolates, and plasmids identified from IMG/PR metagenomes. Hereafter, we refer to these datasets as RefSeq, IMG/PR (I), and IMG/PR (M).

### Mobility of plasmids

The distribution of plasmid mobility types varies notably across the three datasets (Figure 1). In RefSeq, plasmids are nearly evenly distributed among pCONJ (28.6%), pMOB (33.6%), and pNT (37.8%) categories. In contrast, IMG/PR (I) plasmids show a higher prevalence of pMOB (53.2%) with lower proportions of pCONJ (19.0%) and pNT (27.8%). This trend is even more pronounced in IMG/PR (M), where pMOB dominate (67.5%), pCONJ constitute only 3.3% and pNT 29.2%.

**Figure 1.**
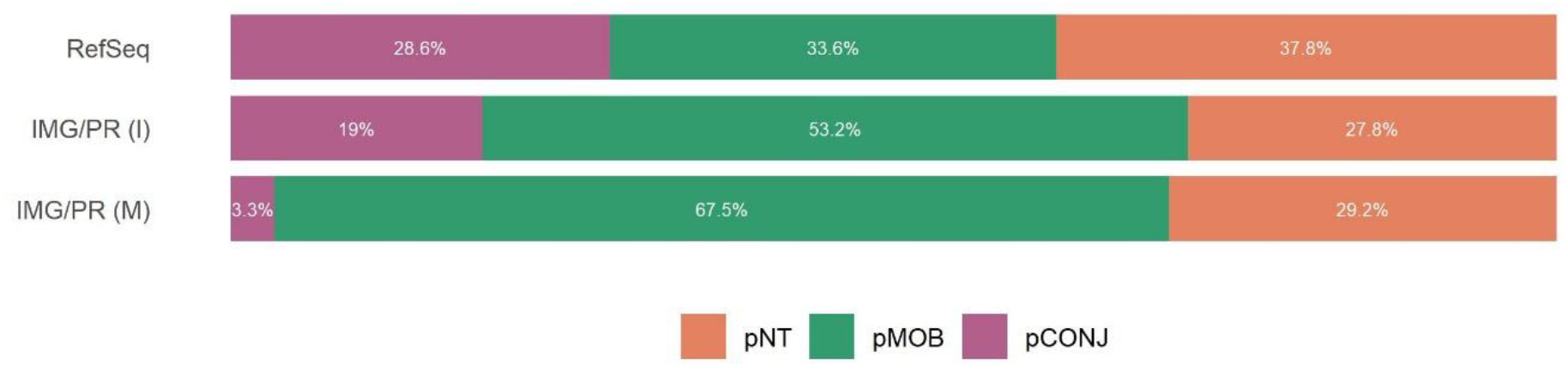
Proportion of plasmid mobility types within each dataset. The proportions of pCONJ, pMOB and pNT among the three datasets are statistically different (Chi-square test, χ^2^(4, N = 189,060) = 26106, p < 2.2×10^−16^, with a medium effect size, Cramér’s V = 0.263, CI_95%_ = [0.260, 0.266]). In RefSeq, pCONJ (adj. res. = 128.3) and pNT (adj. res. = 32.1) are observed more frequently than expected, whereas pMOB is underrepresented (adj. res. = -108.6). In IMG/PR (I), pCONJ occurs more often than expected (adj. res. = 50.0), contrary to pMOB (adj. res. = -20.9) and pNT (adj. res. = -10.4). In IMG/PR (M), pMOB is overrepresented (adj. res. = 105.9), with pCONJ (adj. res. = -143.5) and pNT (adj. res. = -19.3) occurring less frequently than expected. pCONJ are represented in purple, pMOB in green, and pNT in orange.

The median plasmid size differs across the three datasets for all mobility types. It is always highest in RefSeq, followed by IMG/PR (I), and lowest in IMG/PR (M) (Figure 2).

**Figure 2.**
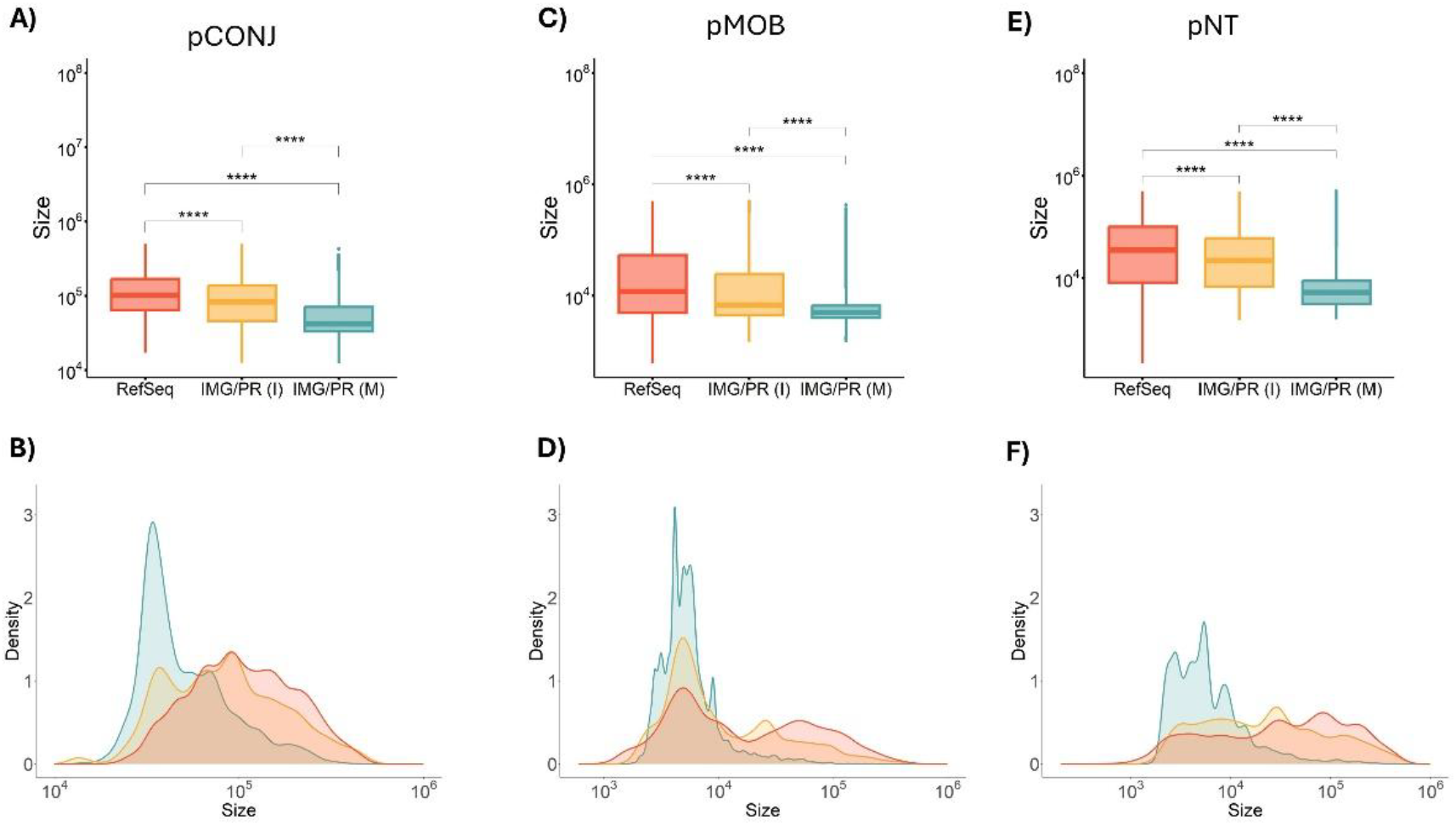
Plasmid size of the three plasmid mobility types across datasets. (A) and (B) pCONJ. Sizes of pCONJ from the three datasets are different (Kruskal-Wallis analyses: χ^2^(2, N = 18,992) = 3365, *p* < 2.2×10^−16^, η^2^ = 0.177, with a large effect size). A Dunn’s test revealed that pCONJ from Refseq are larger than pCONJ from IMG/PR (I) (Z = -18.8, p.adj < 2.2×10^−16^) and from IMG/PR (M) (Z = 58.0, p.adj < 2.2×10^−16^); and that pCONJ from IMG/PR (I) are larger than pCONJ IMG/PR (M) (Z = 34.4, p.adj < 2.2×10^−16^). **(C) and (D) pMOB**. Sizes of pMOB from the three datasets are different (Kruskal-Wallis analyses: χ^2^(2, N = 112,142) = 8843, *p* < 2.2×10^−16^, η^2^ = 0.079, with a moderate effect size). A Dunn’s test revealed that pMOB from Refseq are larger than pMOB from IMG/PR (I) (Z = -18.9, p.adj < 2.2×10^−16^) and from IMG/PR (M) (Z = 80.3, p.adj < 2.2×10^−16^), and that pMOB from IMG/PR (I) are larger than from IMG/PR (M) (Z = 58.6, p.adj < 2.2×10^−16^). **(E) and (F) pNT**. Sizes of pNT from the three datasets are different (Kruskal-Wallis analyses: χ^2^(2, N = 57,926) = 13199, *p* < 2.2×10^−16^, η^2^ = 0.228, with a large effect size). A Dunn’s test revealed that pNT from Refseq are larger than pNT from IMG/PR (Z = -8.2, p.adj < 2.2×10^−16^) and from IMG/PR (M) (Z = 103.2, p.adj < 2.2×10^−16^), and that pNT from IMG/PR (I) are larger than pNT from IMG/PR (M) (Z = 70.0, p.adj < 2.2×10^−16^). Figures showing densities normalize the area under the curve to one for each plasmid type.\**p* < 0.05, \*\**p* < 0.01, and \*\*\**p* < 0.001.

### Linking Antimicrobial Resistance to Plasmid Mobility

To find out whether certain mobility types of plasmid tend to carry more or less ARGs, and to assess whether this analysis may be influenced by the type of plasmid dataset used, we performed a comprehensive screening of plasmids to detect ARGs using the AMRFinderPlus (20) database. Plasmids from RefSeq have the highest prevalence of ARGs (26.4%), followed by those from IMG/PR (I) (15.8%) and IMG/PR (M) (3.1%) (Figure 3A).

**Figure 3A.**
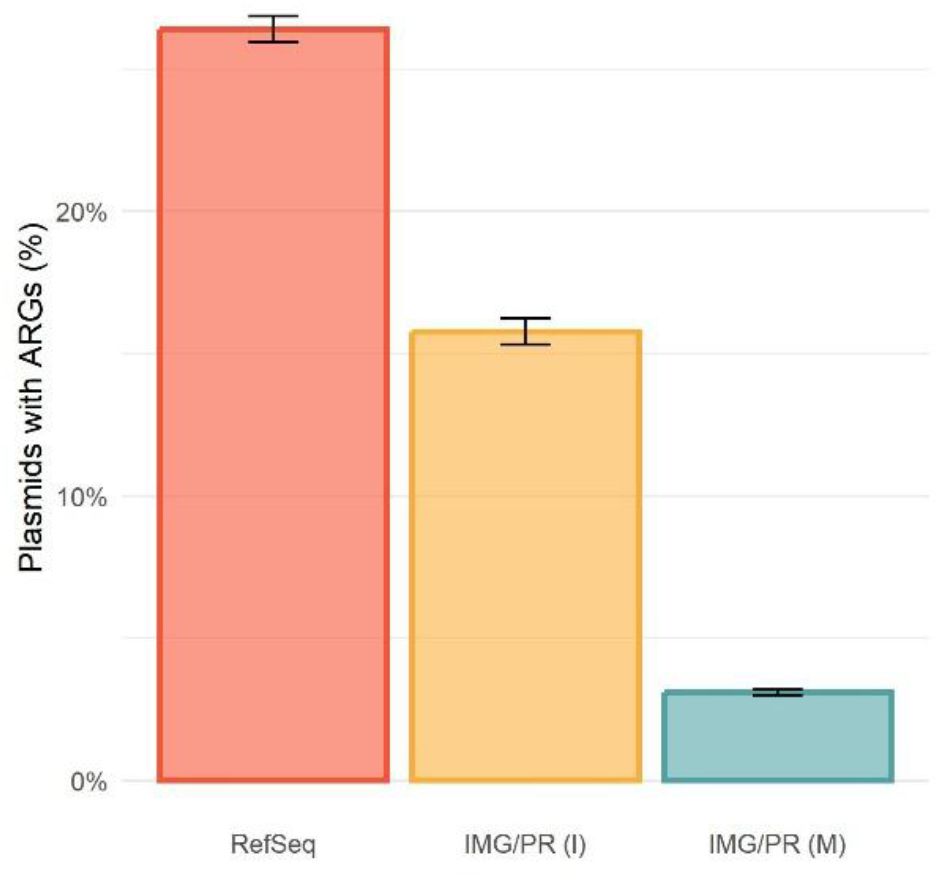
Proportion of plasmids with ARGs in the three datasets. RefSeq contains 9,259 plasmids with ARGs (out of 35,075, hence 26,4% (CI_95%_ = [25,9%, 26,9%]) of the plasmids); IMG/PR (I) contains 3,890 of 24,665 plasmids with ARGs (15.8%, CI_95%_ = [15.3%, 16.2%]), and IMG/PR (M) contains 4,000 of 129,320 plasmids with ARGs (3.1%, CI_95%_ = [3.0%, 3.2%]). The proportion of plasmids with ARGs is different among the three datasets (Chi-square test: *χ*^2^(2) =19712, *p* < 2.2 × 10^−16^, with a large effect size, Cramér’s V = 0.323, CI_95%_ = [0.318, 0.328]). There are more plasmids from RefSeq (adj. res. = 125.2) and IMG/PR (I) (adj. res. = 39.3) with ARGs than expected, but fewer plasmids from IMG/PR (M) with ARGs than expected (adj. res. = -133.2).

The distribution of plasmid mobility types among ARG-carrying plasmids in the three datasets is represented in Figure 3B. **I**n IMG/PR (I), there are more pCONJ and pMOB than expected, and fewer pNT. Among IMG/PR (M) plasmids carrying ARGs, the association between plasmid mobility type and the presence of ARGs is also significant, but the effect size is very small (Figure 3B).

**Figure 3B.**
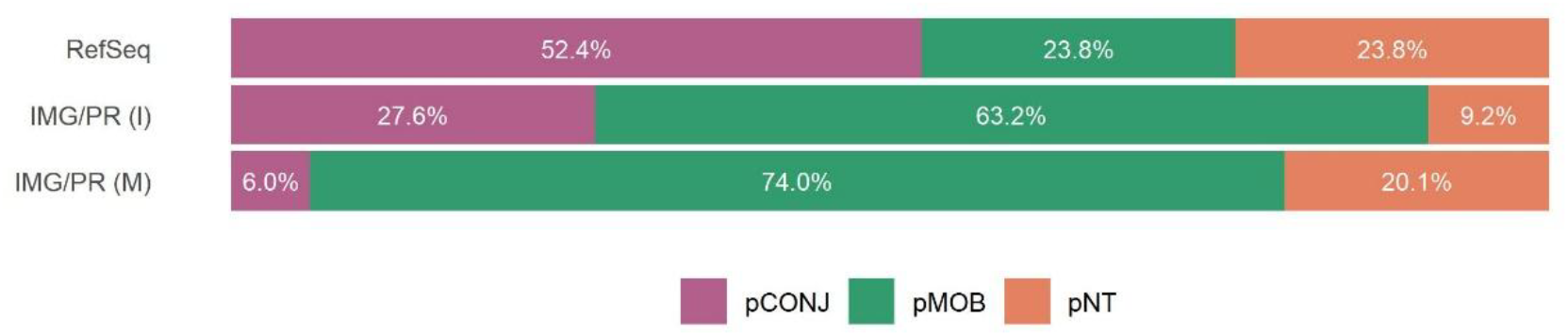
Distribution of plasmid mobility types among ARG-carrying plasmids in the three datasets. For each dataset, only plasmids carrying ARGs were considered. The figure shows the pCONJ, pMOB, and pNT percentages among these ARG-carrying plasmids. Among RefSeq plasmids carrying ARGs, the association between plasmid mobility type and the presence of ARGs is statistically significant (Chi-square test: *χ*^2^(2) = 3490, *p* < 2.2×10^−16^, with a large effect size, Cramér’s V = 0.315, CI_95%_ = [0.304, 0.327]). There are more pCONJ with ARGs than expected (adj. res. = 59.0), but fewer pMOB with ARGs than expected (adj. res. = -23.2), and fewer pNT with ARGs than expected (adj. res. = -32.3). Among IMG/PR (I) plasmids carrying ARGs, the association between plasmid mobility type and the presence of ARGs is also statistically significant (Chi-square test: *χ*^2^(2) = 842, *p* < 2.2×10^−16^, although with a small effect size, Cramér’s V = 0.185, CI_95%_ = [0.176, 0.195]). There are more pCONJ with ARGs than expected (adj. res. = 15.0), and more pMOB with ARGs than expected (adj. res. = 13.6), but fewer pNT with ARGs than expected (adj. res. = -28.2). Among IMG/PR (M) plasmids carrying ARGs, the association between plasmid mobility type and the presence of ARGs is also statistically significant (Chi-square test: *χ*^2^(2) = 234, *p* < 2.2×10^−16^, but with a very small effect size, Cramér’s V = 0.043, CI_95%_ = [0.037, 0.049]). There are more pCONJ with ARGs than expected (adj. res. = 9.6), and more pMOB with ARGs than expected (adj. res. = 8.9), but fewer pNT with ARGs than expected (adj. res. = -13.0). pCONJ are represented in purple, pMOB in green, and pNT in orange.

### Linking Virulence to Plasmid Mobility

We also wanted to assess whether virulence genes are more commonly associated with specific types of plasmid mobility., We have thus analyzed plasmids for the presence of virulence genes (VGs). We observed that, across all three datasets, the percentage of detected virulence genes was low. Only 6.0% of RefSeq plasmids, 4.5% of IMG/PR (I), and 0.09% of IMG/PR (M) plasmids contained VGs (Figure 4A).

**Figure 4A.**
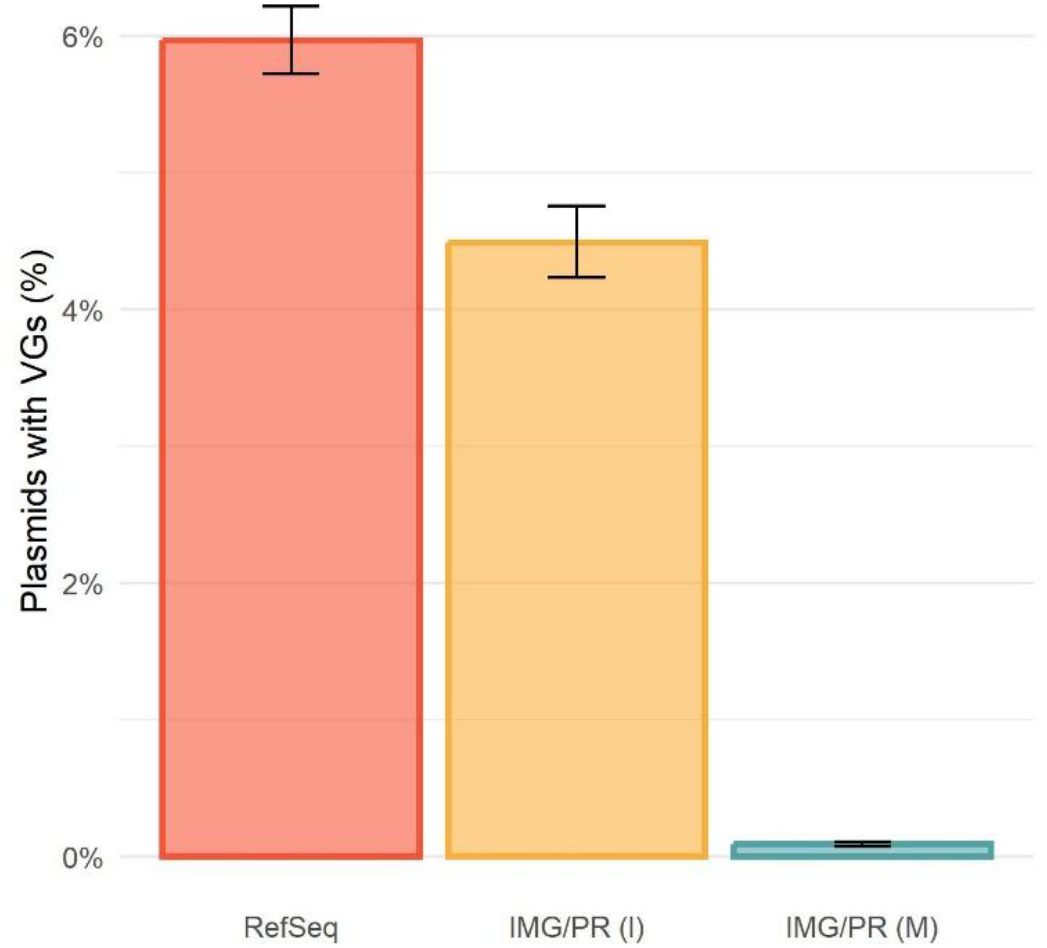
Proportion of plasmids with VGs in the three datasets. RefSeq contains 2,092 plasmids with VGs (out of 35,075, hence 6.0% of the plasmids, CI_95%_ = [5.7%, 6.2%]), IMG/PR (I) contains 1,107 plasmids with VGs (out of 24,665, hence 4.5% of the plasmids, CI_95%_ = [4.2%, 4.8%]), and IMG/PR (M) contains 116 plasmids with VGs (out of 129,320, hence 0.09% of the plasmids, CI_95%_ = [0.07%, 0.11%]). The proportion of plasmids with VGs is different among the three datasets (*χ*^2^(2) =6759, *p* < 2.2 × 10^−16^, with a small effect size, Cramér’s V = 0.189, CI_95%_ = [0.185, 0.193]). There are more plasmids from RefSeq (adj. res. = 66.6) and from IMG/PR (I) (adj. res. = 35.1) with VGs than expected, but fewer plasmids from IMG/PR (M) with VGs than expected (adj. res. = -81.1).

The distribution of plasmid mobility types among VG-carrying plasmids in the three datasets is represented in Figure 4B. Among RefSeq plasmids, there are more pCONJ and pNT than expected, and fewer pMOB than expected. In IMG/PR (I) and IMG/PR (M), pCONJ are found more often than expected, while pMOB and pNT are less frequent than expected (Figure 4B).

**Figure 4B.**
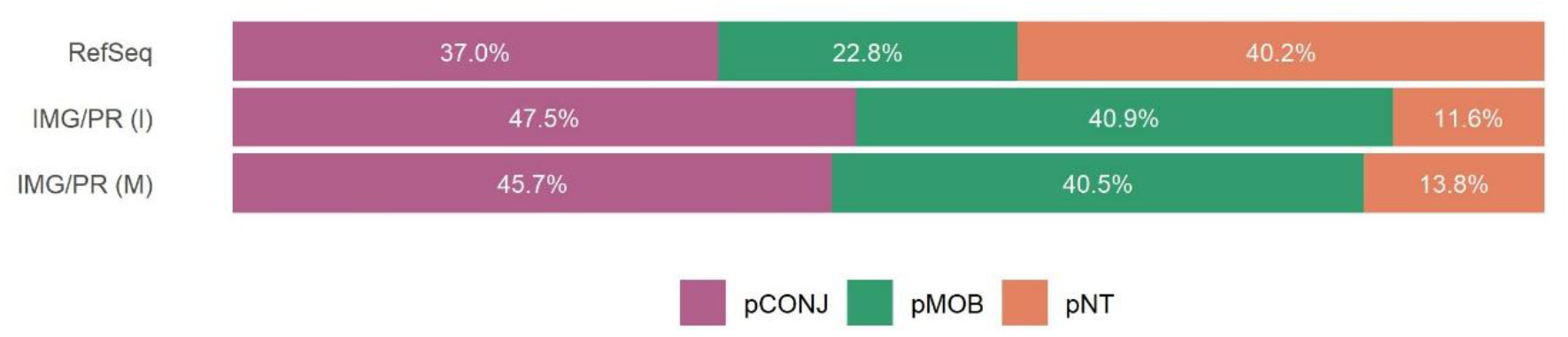
Distribution of plasmid mobility types among VG-carrying plasmids in the three datasets. For each dataset, only plasmids carrying VGs were considered. The figure shows the pCONJ, pMOB, and pNT percentages among these VG-carrying plasmids. Among RefSeq plasmids carrying VGs, the association between plasmid mobility type and the presence of VGs is statistically significant (χ^2^(2) = 134, p < 2.2×10-16, although with a very small effect size, Cramér’s V = 0.062, CI95% = [0.051, 0.072]). There are more pCONJ (adj. res. = 8.7) and pNT (adj. res. = 2.3) with VGs than expected, but fewer pMOB with VGs than expected (adj. res. = -10.7). Among IMG/PR (I) plasmids carrying VGs, 47.5% are pCONJ, 40.9% are pMOB, and 11.6% are pNT. In this dataset, the association between plasmid mobility type and the presence of VGs is also statistically significant (χ^2^(2) = 640, p < 2.2×10-16, and also with a small effect size, Cramér’s V = 0.161, CI95% = [0.146, 0.177]). There are more pCONJ with VGs than expected (adj. res. = 24.8), but fewer pMOB (adj. res. = -8.4) and pNT (adj. res. = -12.3) with VGs than expected. Among IMG/PR (M) carrying VGs, 45.7% are pCONJ, 40.5% are pMOB, and 13.8% are pNT. In this dataset, the association between plasmid mobility type and the presence of VGs is also statistically significant (χ^2^(2) = 654, p < 2.2×10-16, and also with a very small effect size, Cramér’s V = 0.071, CI95% = [0.054, 0.089]). There are more pCONJ with VGs than expected (adj. res. = 25.6), but fewer pMOB (adj. res. = -6.2) and pNT (adj. res. = -3.7) with VGs than expected. pCONJ are represented in purple, pMOB in green, and pNT in orange.

### Taxonomic Composition of Plasmid Hosts

The host organisms represented in each dataset may influence the presence of resistance and virulence genes, as well as plasmid mobility types. To explore this, we assessed plasmid hosts’ taxonomic diversity and distribution. For taxonomic analyses, we considered only plasmids with an identified genus. Genus-level host information is available for all plasmids in RefSeq, but only for 97.4% of the plasmids (24,022) in IMG/PR (I), and 21.4 % of the plasmids (24,708) in IMG/PR (M). We selected 30 genera with over 1% representation in at least one dataset.

In RefSeq, the three most prevalent genera are *Escherichia* (17.2%), *Klebsiella* (13.8%), and *Enterococcus* (4.9%), whereas, in IMG/PR (I), *Escherichia* (16.0%), *Staphylococcus* (9.3%), and *Salmonella* (4.2%) are the most common. In IMG/PR (M), *Bifidobacterium* (10.0%), *Staphylococcus* (9.9%), and *Escherichia* (7.7%) are the most represented hosts (Figure 5; Supplementary file “genus_dataset_percentage.xlsx”). Therefore, *Escherichia* is consistently one of the most prevalent genera across all three datasets.

**Figure 5.**
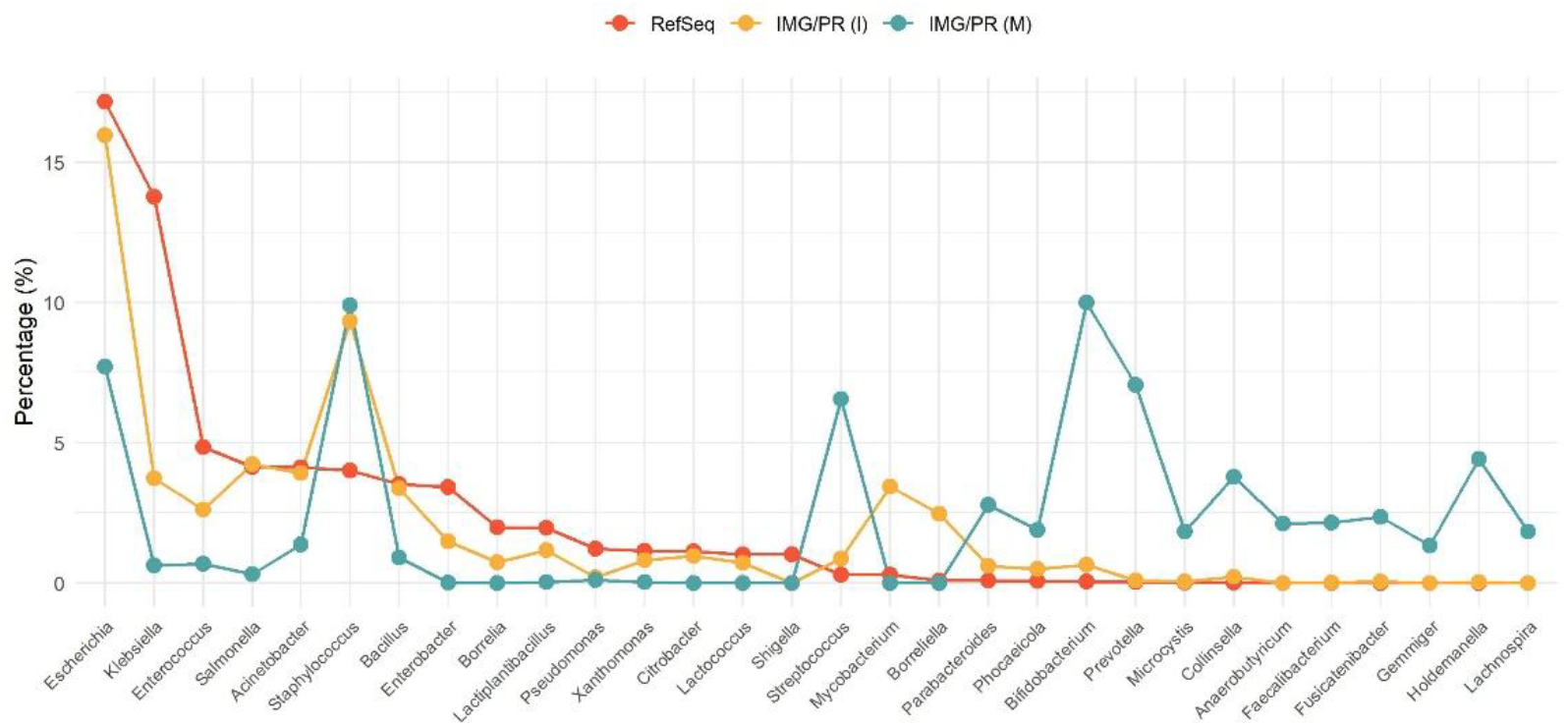
Genus-level distribution of plasmid hosts across datasets. Relative abundance (%) of the 30 most represented genera among plasmid hosts in each dataset. Only plasmids with an assigned host genus were included, and only genera with >1% representation in at least one dataset are shown. The x-axis displays genera; the y-axis indicates the percentage of plasmids associated with each genus. Red, yellow, and blue lines represent RefSeq, IMG/PR (I), and IMG/PR (M), respectively.

Among the 30 most represented genera, the RefSeq and IMG/PR (I) profiles are broadly similar regarding genus-level composition. *Staphylococcus, Mycobacterium*, and *Borreliella* are more abundant in IMG/PR (I) (Figure 5).

Compared to RefSeq and IMG/PR (I), plasmids from IMG/PR (M) show a distinct taxonomic profile, with higher representation of genera such as *Streptococcus, Bifidobacterium, Prevotella*, and *Collinsella* (Figure 5). Additionally, *Lachnospira, Gemmiger*, and *Anaerobutyricum* were exclusively detected in IMG/PR (M).

Plamids from *Holdemanella* were present in both IMG/PR (M) and IMG/PR (I) but absent from RefSeq. In contrast, genera such as *Borrelia, Borreliella, Citrobacter, Mycobacterium*, and *Lactococcus* were found only in the isolate datasets and not in metagenomic data. *Shigella* was detected exclusively in RefSeq.

Overall, genus-level profiles from the two isolate-based datasets are more similar to each other than to the metagenome-based dataset, suggesting that host taxonomic composition of the datasets differs substantially depending on whether they are isolate or metagenomic based.

### Linking Taxonomy and Antimicrobial Resistance

Among plasmids with assigned host genus, 1,612 (2.5%) plasmids from IMG/PR (M) and 3,724 (15.5%) plasmids from IMG/PR (I) carry ARGs. As previously noted, 9,259 (26.4%) plasmids in RefSeq harbor ARGs (all have genus classification). When considering only the plasmids carrying ARGs, 86.1% of those in IMG/PR (M) belong to the genus *Staphylococcus*, 3.5% to *Klebsiella*, and less than 1% to *Escherichia* (0.7%). In IMG/PR (I), the majority of ARG-carrying plasmids are also from *Staphylococcus* (38.9%), followed by *Escherichia* (13.9%), *Klebsiella* (12.2%), and *Acinetobacter* (6.0%). However, in RefSeq, *Staphylococcus* ranks only fourth among genera with ARG-carrying plasmids (6.9%). The top three genera with the highest proportions of ARG-carrying plasmids in RefSeq are *Klebsiella* (26.8%), *Escherichia* (24.8%), and *Enterococcus* (7.1%).

Given the observed differences in the genera contributing to antimicrobial resistance across the three datasets, we questioned whether the higher prevalence of ARGs in RefSeq is due to a higher representation of specific genera with more ARGs, or if resistance levels are elevated across all genera in this dataset. In other words, if genus *X* shows high ARG prevalence consistently in all datasets but is more abundant in RefSeq, the higher resistance in RefSeq could be explained simply by its taxonomic composition. Conversely, if genera *X, Y*, and *Z* all exhibit greater ARG prevalence in RefSeq compared to the other datasets, this would suggest that the RefSeq dataset is enriched in ARGs irrespective of host genus composition.

When analyzing the most abundant genera in RefSeq — namely *Acinetobacter, Enterobacter, Enterococcus, Escherichia, Klebsiella, Salmonella*, and *Staphylococcus*—we observe that the percentage of plasmids carrying ARGs is consistently higher in RefSeq for all genera except

*Staphylococcus*. For *Staphylococcus*, plasmids from IMG/PR (I) show the highest ARG prevalence, followed by the plasmids from IMG/PR (M), with plasmids from RefSeq s having the lowest proportion (Table 1). These results indicate that the overall enrichment of ARGs in plasmids from RefSeq is not solely due to one or two highly resistant genera. They rather reflect a broader trend of higher resistance gene prevalence across most genera in this dataset.

**Table 1.**
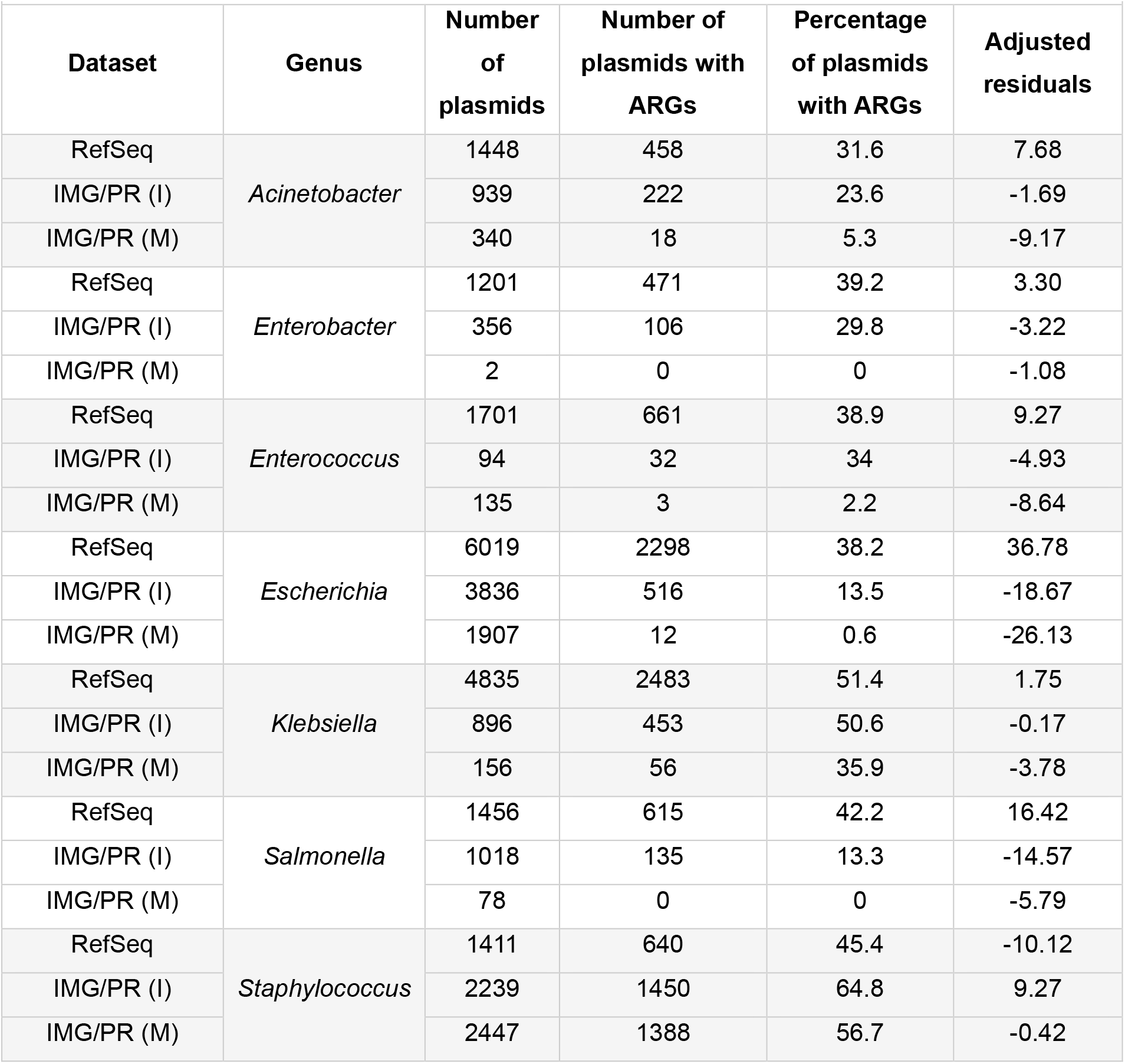
Proportion of plasmids carrying ARGs in each dataset, by genus. Adjusted residuals were calculated using Chi-squared test.

**Table 2.** Distribution of plasmid mobility types across datasets within each genus.

| Genus | Dataset | N | pCON<br>J<br>(%) | Adjusted<br>Residuals<br>(pCONJ) | pMOB<br>(%) | Adjusted<br>Residuals<br>(pMOB) | pNT<br>(%) | Adjusted<br>Residuals<br>(pNT) |
| --- | --- | --- | --- | --- | --- | --- | --- | --- |
| *Acinetobacter* | RefSeq | 1448 | 13.8 | 0.64 | 34.4 | -7.87 | 51.8 | 7.35 |
|  | IMG/PR (I) | 939 | 13.8 | 0.47 | 53.2 | 9.13 | 32.9 | -9.36 |
|  | IMG/PR (M) | 340 | 10.6 | -1.64 | 38.2 | -1.25 | 51.2 | 2.36 |
| *Enterobacter* | RefSeq | 1201 | 37.4 | -0.11 | 39 | -5.39 | 23.6 | 6.82 |
|  | IMG/PR (I) | 356 | 37.9 | 0.20 | 55.3 | 5.51 | 6.7 | -7.07 |
|  | IMG/PR (M) | 2 | 0 | -1.10 | 0 | -1.22 | 100 | 2.84 |
| *Enterococcus* | RefSeq | 1701 | 18.6 | 2.05 | 41.4 | -12.3 | 40.0 | 11.53 |
|  | IMG/PR (I) | 94 | 52.1 | -3.41 | 35.1 | 9.63 | 12.8 | -7.51 |
|  | IMG/PR (M) | 135 | 29.6 | 2.07 | 65.9 | 6.30 | 4.4 | -8.40 |
| *Escherichia* | RefSeq | 6019 | 40.7 | 31.26 | 42.1 | -42.02 | 17.2 | 20.45 |
|  | IMG/PR (I) | 3836 | 20.4 | -12.08 | 74.3 | 21.12 | 5.3 | -14.32 |
|  | IMG/PR (M) | 1907 | 3.5 | -26.05 | 91.5 | 30.13 | 5.0 | -9.52 |
| *Klebsiella* | RefSeq | 4835 | 39.4 | -1.80 | 29.9 | -5.76 | 30.7 | 7.88 |
|  | IMG/PR (I) | 896 | 41.3 | 0.90 | 40.2 | 6.08 | 18.5 | -7.23 |
|  | IMG/PR (M) | 156 | 48.7 | 2.27 | 32.1 | 0.15 | 19.2 | -2.62 |
| *Salmonella* | RefSeq | 1456 | 50.0 | 0.64 | 42.9 | -2.49 | 7.1 | 4.01 |
|  | IMG/PR (I) | 1018 | 51.4 | 1.58 | 46.3 | 1.06 | 2.4 | -5.76 |
|  | IMG/PR (M) | 78 | 14.1 | -6.34 | 67.9 | 4.14 | 17.9 | 4.85 |
| *Staphylococcus* | RefSeq | 1411 | 3.8 | 3.61 | 36.5 | -26.72 | 59.7 | 26.04 |
|  | IMG/PR (I) | 2239 | 3.3 | 3.20 | 80.8 | 18.49 | 15.9 | -19.93 |
|  | IMG/PR (M) | 2447 | 1.0 | -6.25 | 69.6 | 4.81 | 29.4 | -2.80 |

### Linking Taxonomy and Plasmid Mobility

We previously observed that the frequency of the different plasmid mobility types varies across the three datasets. This raises the question of whether these differences are driven by the relative abundance of plasmids from each genus within each dataset or if plasmid mobility patterns differ within the same genera across the datasets.

In most genera analyzed (5 out of 7), the proportion of pCONJ is similar between RefSeq and IMG/PR (I). However, for *Escherichia* — the most abundant genus in both RefSeq (17.2%) and IMG/PR (I) (16.0%) — the proportion of pCONJ plasmids is substantially higher in RefSeq (40.7%) compared to IMG/PR (I) (20.4%). Therefore, the overall differences in pCONJ proportions between datasets may be driven by the presence and relative abundance of specific genera, such as *Escherichia*. In other words, the high representation of this genus in the datasets could explain why the global pCONJ proportion is greater in RefSeq (28.6%) than in IMG/PR (I) (19.0%). Additionally, in 5 out of the 7 genera, the proportion of pCONJ is higher in both isolate-derived datasets (RefSeq and IMG/PR (I)) than in the IMG/PR (M), a bias that may reflect the differences in the efficacy of assembling plasmids from isolates and from metagenomes. This suggests that conjugation-related mechanisms may be underdetected or more difficult to identify in plasmids assembled from metagenomes. Interestingly, for *Enterococcus* and *Klebsiella*, the proportion of pCONJ plasmids is higher in both IMG/PR datasets than in RefSeq.

For pMOB, 7 out of 8 genera show a higher proportion in IMG/PR (I) than in RefSeq. In contrast, for pNT, all genera exhibit higher proportions in RefSeq compared to IMG/PR (I). This pattern is consistent with what is observed when analysing the full datasets. Therefore, the proportion of pMOB and pCONJ does not seem to be influenced by the presence or abundance of any specific genus within the dataset.

### Putative bias in metagenomic data

One limitation of studying plasmids assembled from metagenomes is that it favours small plasmids. Because of their high copy number and simpler structure, these plasmids are often more easily recovered in metagenomic assemblies. In contrast, large plasmids, which often contain repetitive and complex regions, are more prone to misassembly — frequently breaking into multiple fragments that may be mistakenly identified as several smaller, false plasmids. Therefore, we analysed the percentage of plasmids with resistance genes by removing different slices of the smallest plasmids in the IMG/PR (M) dataset. Naturally, this changes the proportions of each plasmid mobility type in the sample (Table 1). Considering only the 30% largest plasmids, the percentage of pCONJ increases from 3.30% to 11.01%. While the percentage of pNT also increases, the percentage of pMOB decreases (Table 1).

**Table 1.** Percentage of each plasmid mobility type, removing different slices of the smaller plasmids in the IMG/PR (M) dataset.

| % plasmids considered | Nr plasmids considered | Mean length | Min length | pCONJ (%) | pMOB (%) | pNT (%) |
| --- | --- | --- | --- | --- | --- | --- |
| 100 | 129,320 | 9996.8 | 1,546 | 3.30 | 67.45 | 29.25 |
| 90 | 116,388 | 10831.1 | 2,834 | 3.67 | 69.63 | 26.70 |
| 80 | 103,456 | 11795.0 | 3,463 | 4.13 | 69.67 | 26.20 |
| 70 | 90,524 | 12933.2 | 4,138 | 4.72 | 69.11 | 26.17 |
| 60 | 77,592 | 14375.3 | 4,551 | 5.50 | 66.59 | 27.91 |
| 50 | 64,660 | 16275.3 | 5,165 | 6.60 | 63.63 | 29.77 |
| 40 | 51,728 | 18979.9 | 5,697 | 8.25 | 62.67 | 29.07 |
| 30 | 38,796 | 23270.7 | 6,693 | 11.01 | 55.21 | 33.78 |

However, this analysis caused practically no change in the percentage of plasmids with ARGs since we observed 3.09% when the whole sample was considered and 5.53% when removing 70% of the smallest plasmids (Table 2).

**Table 2.**
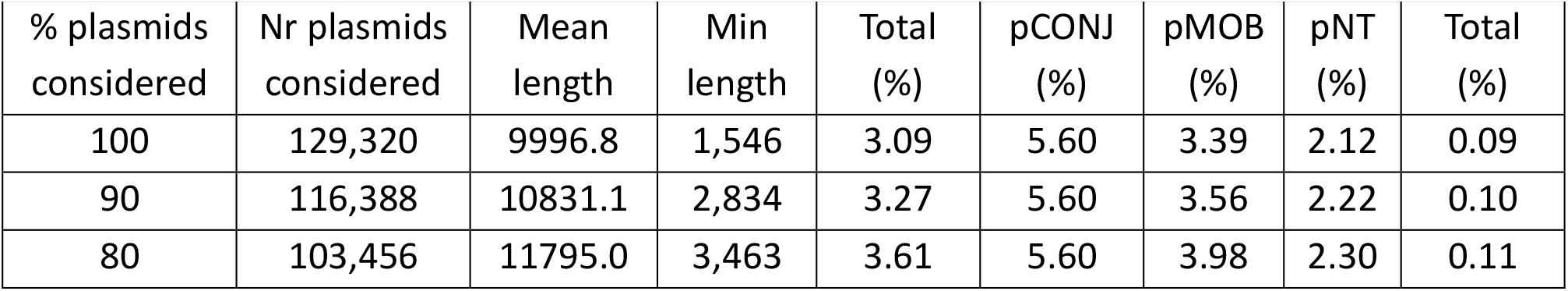
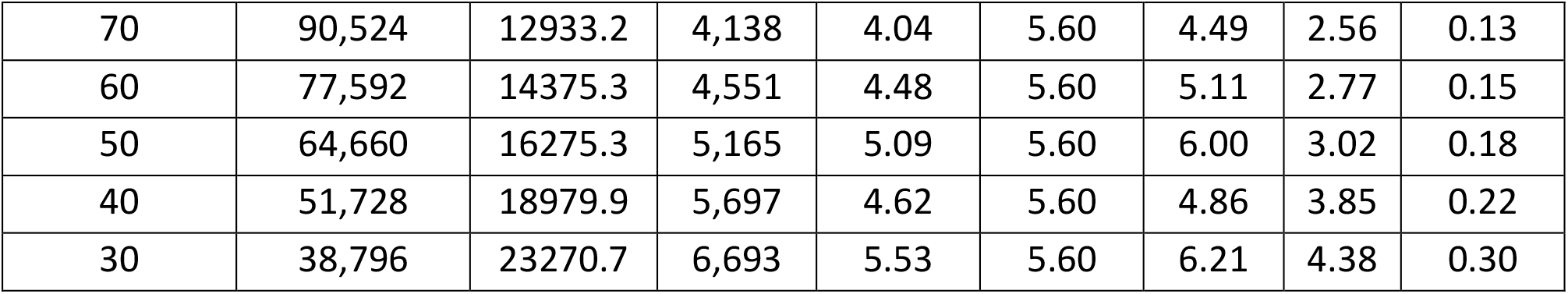
Percentage of each plasmid mobility type carrying ARGs, removing different slices of the smaller plasmids in the IMG/PR (M) dataset.

Different identity thresholds are used to detect resistance and virulence genes in samples from isolates and metagenomes. To account for this, we varied the identity thresholds in our analyses regarding plasmids assembled from metagenomes. Our findings indicate that these variations had minimal impact on the proportions of ARGs. We identified ARGs in 3.52% of plasmids with 70% identity and 3.09% with 90% identity. Additionally, the percentage of resistance detected remains relatively consistent across different plasmid mobility types (Table 3).

**Table 3.** Percentage of each plasmid mobility type carrying ARGs considering different identity thresholds in the IMG/PR (M) dataset.

| % identity | Total (%) | pCONJ (%) | pMOB (%) | pNT (%) |
| --- | --- | --- | --- | --- |
| 90 | 3.09 | 5.60 | 3.39 | 2.12 |
| 80 | 3.45 | 6.00 | 3.89 | 2.16 |
| 70 | 3.52 | 6.14 | 3.98 | 2.20 |

## Discussion

This work aimed to evaluate how the use of different datasets of plasmids could influence the analysis, and conclusions drawn. We analysed a comprehensive collection of fully sequenced plasmids derived from isolates, IMG/PR (I), and metagenomic samples, IMG/PR (M), from IMG/PR database (21), and plasmids from a non-redundant and curated database, the NCBI Reference Sequence Database (RefSeq) (18).

Our comparative analysis of these three plasmid datasets reveals significant differences in key plasmid features, including antimicrobial resistance, virulence, mobility type, and host taxonomy. Even though we selected only putative complete plasmids in IMG/PR (M), their median sizes across all plasmid mobility types are smaller compared to those in RefSeq and IMG/PR (I) (Figure 2). This suggests that, despite filtering for plasmid completeness, metagenomic assembly may still favour the recovery of shorter plasmids, maybe because longer plasmids are more challenging to assemble in complex microbial communities.

We observed differences in the distribution of plasmid mobility types across datasets (Figure 1). Compared to the other datasets, plasmids from the RefSeq are enriched in pCONJ, which play a key role in horizontal gene transfer and resistance dissemination. This result may be explained by the nature of this dataset which is likely to be enriched with bacterial pathogens.

On the other hand, IMG/PR (M) exhibits a strong predominance of pMOB and an underrepresentation of pCONJ. Could the lower number of conjugative plasmids in IMG/PR(M) be due to the fragmentation of large plasmids into smaller pieces during sequencing and assembly, leading to an overrepresentation of false small plasmids? Our results suggest that this may be the case. We eliminated a large proportion of the smaller plasmids from IMG/PR (M). Considering only the largest 30% of plasmids, we observed that the percentage of pCONJ is 11.01% (Table 1). However, both this value and the percentage of pCONJ observed in IMG/PR (I) (19.0%, Figure 1) are lower than the values observed in RefSeq (28.6%) and from previous estimates reported in other works (approximately 25%) (2, 3, 10, 17).

While conjugative plasmids can diversify more frequently than less mobile ones by acquiring DNA from other mobile genetic elements and host chromosomes ^20–22^, three key factors suggest their proportion should be lower. First, with a few exceptions (22–24), several studies suggested that the transfer rate of pCONJ is too low to compensate for its fitness cost (25). Secondly, if plasmid transfer imposes a cost on bacteria, pCONJ are likely more costly than pMOB, which, in turn, should be more costly than pNT (26). Third, pCONJ may suffer deactivating mutations (i.e., any DNA changes, such as SNPs, or DNA rearrangements) in transfer genes, making them pMOB or pNT (3, 9–12, 15, 27). Recent studies corroborate this view. Phylogenetic analyses of relaxases indicate more transitions from pCONJ to pMOB than the reverse (17). Pointing in the same direction, a recent analysis of pseudogenes in enterobacterial plasmids shows that the non-functionalization of the conjugation machinery has led to the emergence of pMOB (28). According to these mutational events, the frequency of pCONJ is expected to decrease in favor of pMOB or pNT. Likewise, through mutational events, the frequency of pMOB may decrease in favor of pNT (28).

Regarding antimicrobial resistance, RefSeq plasmids have the highest proportion of ARGs (26.4%), followed by IMG/PR (I) (15.8%) and IMG/PR (M) (3.1%). These statistically significant differences indicate that ARGs are more frequently detected in plasmids from isolate-based datasets than metagenomic sources. The elevated ARG prevalence in RefSeq may reflect a sampling bias toward bacterial genomes from clinically relevant strains subjected to higher selective pressures to maintain antimicrobial resistance. Our results confirm that plasmids exhibit a similar trend of having a higher percentage of resistance, emphasizing their pivotal role in driving genome evolution under antibiotic selection pressure. In contrast, the lower ARG content in IMG/PR (M) could be attributed to biological factors—such as a greater representation of environmental or commensal bacteria—and technical limitations in metagenomic assembly and annotation, which may hinder the detection of resistance determinants.

An interesting pattern emerges with *Escherichia*, which is abundant in all datasets but shows very different levels of ARGs (Table 1). While in the RefSeq dataset, *Escherichia* plasmids have a high proportion of resistance genes (38.2%), the percentage in IMG/PR (I) is lower (13.5%), and IMG/PR (M) *Escherichia* plasmids rarely carry ARGs (0.6%). Again, this difference could reflect the nature of the samples: RefSeq likely includes more clinical isolates under stronger antibiotic pressure, leading to higher ARG prevalence, whereas metagenomic samples may represent more diverse or environmental populations where *Escherichia* has less selective pressure for antibiotic resistance. This discrepancy highlights how sampling can shape the apparent resistance profiles even within the same genus, making *Escherichia* an excellent example of how dataset origin impacts results and their interpretations. When we test the association between antibiotic resistance and plasmid mobility type, we observe that pCONJ are always statistically significant above expected in all datasets, suggesting that antimicrobial resistance is linked to conjugation (Figure 3B).

Our analyses of robustness considered different identity thresholds (70%, 80%, and 90%) for detecting resistance genes in metagenomic data (Table 3). Using different thresholds of identity in metagenomic studies to identify ARGs is essential because metagenomic samples involve complex mixtures of organisms with significant sequence diversity, requiring lower thresholds to capture a broader range of gene variants.

Regarding virulence, we detected a higher percentage of VGs in RefSeq (6.0%), followed by IMG/PR (I) (4.5%) and IMG/PR (M) (0.09%). However, despite the differences among the three datasets, the number of plasmids with identified VGs was very low across all datasets (Figure 4A). This may reflect a true lower prevalence of VGs in plasmids, but it could also be due to limitations in annotation. For instance, some virulence genes may be misclassified or not even identified, particularly when they share similarities with other systems, such as those belonging to the conjugation machinery. Given the small number of detected VGs and the potential for misannotation, we believe conclusions regarding virulence genes should be interpreted cautiously, as their identification may be more prone to errors or ambiguity.

Host taxonomic analysis allows us to further explore the differences between datasets. The RefSeq and IMG/PR (I) datasets have similar genus-level compositions, and they are different from the taxonomic distribution of IMG/PR (M) (Figure 5). Some genera are uniquely found in either isolate-based or metagenomic datasets, illustrating how sampling shapes the taxonomic diversity of plasmids. These taxonomic differences may influence plasmid analyses, especially when specific taxa dominate datasets.

Our findings indicate that the elevated ARG prevalence in RefSeq is not solely driven by the overrepresentation of a few highly resistant genera. When examining the most abundant genera across datasets—such as *Escherichia, Klebsiella, Enterococcus*, and *Acinetobacter*—we found that RefSeq consistently contains a higher proportion of ARG-carrying plasmids for nearly all of them, except for *Staphylococcus*. IMG/PR (I) and IMG/PR (M) show higher ARG prevalence in this genus. This enrichment of ARGs across genera in RefSeq implies a bias that favors ARG-rich plasmids.

Regarding plasmid mobility, the prevalence of pCONJ appears to be influenced by the taxonomy of the plasmids in the datasets. This is evidenced by the similar percentage of pCONJ across all genera (the seven most frequent in RefSeq), except for *Escherichia* (Table 2). Although there are many plasmids from this genus in all datasets, the overall percentage of pCONJ in the datasets is affected by the individual percentage of pCONJ in *Escherichia* (Table 2). In IMG/PR (M), the lower proportion of pCONJ across all genera suggests possible issues with identifying conjugation mechanisms. The consistent patterns for pMOB — more common in IMG/PR (I) — and pNT — more frequent in RefSeq — across genera, mirror what is observed in the total datasets, suggesting that these types of mobility are more influenced by sample source than by taxonomy.

Our results show significant differences in plasmid mobility types, abundance of resistance and virulence genes across the three distinct datasets analyzed. Although assembling plasmids from metagenomes avoids the selective pressures associated with strain isolation and provides a more representative distribution, it may also lead to an increased risk of false negatives by the assembly of truncated molecules (29) and still fail to capture the full diversity of the plasmidome. This highlights the need for caution when drawing conclusions based on specific datasets, as the sample type and dataset content can strongly influence the outcomes.

## Methods

Plasmid sequences from IMG/PR database were obtained on December 19, 2023 (https://genome.jgi.doe.gov/portal/IMG_PR/IMG_PR.home.html) (21). We selected plasmids classified as putatively complete and removed all plasmids larger than 500 kb that could represent potential secondary chromosomes. We retained plasmids identified across isolate genomes (24,665 sequences) and metagenomes (129,320 sequences), resulting in 153,985 plasmids.

Antibiotic resistance genes (ARGs) were detected using AMRFinderPlus version 4.0.19, database version 2025-03-25.1, with default parameters (20). Virulence genes (VGs) were detected using ABRicate against the Virulence Factors Database (VFDB) (parameters: ‘--db vfdb’), which includes 4,360 sequences (30). VFDB was last updated on 1 February 2024. The AMRFinderPlus identifies resistance genes with a default value of 90% minimum identity. To analyze the robustness of the results in the metagenomic data (IMG/PR (M) dataset), we varied the identity values to 70% and 80%.

Plasmid data from RefSeq (18), which already include information on resistance and virulence genes, were obtained from Coluzzi & Rocha (31). The authors classified the plasmids according to their mobility type using CONJScan (32). Plasmids are categorized as pCONJ, pMOB, pNT, PP (phage-plasmids), and pOriT. We did not consider PP plasmids and treated pOriT as pMOB.

The IMG/PR database (21) contains metadata regarding the presence of mobility genes, type IV coupling protein (T4CP), type IV secretion systems (T4SS), mating pair formation (MPF), obtained using CONJscan HMM models (32), and origin of transfer (*oriT*) obtained against a database of *oriT* sequences (3). We classified plasmids as conjugative (pCONJ), mobilizable (pMOB), or non-transmissible (pNT) based on the machinery required for DNA transfer. Plasmids encoding mobility genes, T4CP, T4SS, and MPF system were classified as pCONJ. If plasmids were not included in the pCONJ category, they were classified as pMOB if they contained mobility genes or an *oriT*. The remaining plasmids were pNT. Nevertheless, these non-transmissible plasmids may have an unidentified *oriT*, owing to the limited understanding of these regions (3).

### Statistics

We performed the statistical analyses in R, version 4.4.2 (33). We used Pearson’s chi-squared test to analyze the dependence between two categories and the respective residuals, measuring effect size using Cramér’s V from rcompanion package, version 2.4.36. We performed a z-test to compare the plasmid proportions and Cohen’s h to measure the effect size. We used the nonparametric Mann-Whitney U test, also known as the Wilcoxon rank-sum test, to compare the means of two samples, and calculate the effect size using the Glass rank biserial correlation coefficient (r) from effectsize package version 1.0.0. We used Dunn’s post-hoc test from FSA package version 0.9.5 to identify which groups differ. Visualizations were performed with the R package ggplot2 version v3.5.1.

## Acknowledgments

The authors thank Eduardo P. C. Rocha and João A. for their critical review of the manuscript and constructive suggestions. Célia P. F. Domingues and João S. Rebelo acknowledge FCT-Fundação para a Ciência e a Tecnologia, IP for their fellowships (PhD grants UI/BD/153078/2022, and SFRH/BD/04631/2021, respectively). FCT also supports cE3c by contract DOI 10.54499/UIDP/00329/2020. The funder had no role in study design, data collection and interpretation, or the decision to submit the work for publication.

## Author contributions

Conceptualization, F.D and T.N.; Software, C.P.F.D. and J.S.R.; Methodology, C.P.F.D., J.S.R., F.D. and T.N.; Investigation, C.P.F.D. and J.S.R.; Formal Analysis, C.P.F.D., J.S.R., F.D. and T.N.; Writing – Original Draft, C.P.F.D., J.S.R., F.D. and T.N..; Resources, T.N.; Visualization, C.P.F.D. and J.S.R.; Supervision, F.D. and T.N.

